# The *Candida albicans* reference strain SC5314 contains a rare, dominant allele of the transcription factor Rob1 that modulates biofilm formation and oral commensalism

**DOI:** 10.1101/2023.06.17.545405

**Authors:** Virginia E. Glazier, Juraj Kramara, Tomye Ollinger, Norma V. Solis, Robert Zarnowski, Rohan S. Wakade, Min-Ju Kim, Gabriel J. Weigel, Shen-Huan Liang, Richard J. Bennett, Melanie Wellington, David R. Andes, Mark A. Stamnes, Scott G. Filler, Damian J. Krysan

## Abstract

*Candida albicans* is a diploid human fungal pathogen that displays significant genomic and phenotypic heterogeneity over a range of virulence traits and in the context of a variety of environmental niches. Here, we show that the effects of Rob1 on biofilm and filamentation virulence traits is dependent on both the specific environmental condition and the clinical strain of *C. albicans*. The *C. albicans* reference strain SC5314 is a *ROB1* heterozygote with two alleles that differ by a single nucleotide polymorphism at position 946 resulting in a serine or proline containing isoform. An analysis of 224 sequenced *C. albicans* genomes indicates that SC5314 is the only *ROB1* heterozygote documented to date and that the dominant allele contains a proline at position 946. Remarkably, the *ROB1* alleles are functionally distinct and the rare *ROB1^946S^*allele supports increased filamentation in vitro and increased biofilm formation in vitro and in vivo, suggesting it is a phenotypic gain-of-function allele. SC5314 is amongst the most highly filamentous and invasive strains characterized to date. Introduction of the *ROB1^946S^*allele into a poorly filamenting clinical isolate increases filamentation and conversion of an SC5314 laboratory strain to a *ROB1^946S^* homozygote increases in vitro filamentation and biofilm formation. In a mouse model of oropharyngeal infection, the predominant *ROB1^946P^* allele establishes a commensal state while the *ROB1^946S^* phenocopies the parent strain and invades into the mucosae. These observations provide an explanation for the distinct phenotypes of SC5314 and highlight the role of heterozygosity as a driver of *C. albicans* phenotypic heterogeneity.

**Importance:** *Candida albicans* is a commensal fungus that colonizes human oral cavity and gastrointestinal tracts but also causes mucosal as well as invasive disease. The expression of virulence traits in *C. albicans* clinical isolates is heterogenous and the genetic basis of this heterogeneity is of high interest. The *C. albicans* reference strain SC5314 is highly invasive and expresses robust filamentation and biofilm formation relative to many other clinical isolates. Here, we show that SC5314 derivatives are heterozygous for the transcription factor Rob1 and contain an allele with a rare gain-of-function SNP that drives filamentation, biofilm formation, and virulence in a model of oropharyngeal candidiasis. These finding explain, in part, the outlier phenotype of the reference strain and highlight the role of heterozygosity plays in the strain-to-strain variation of diploid fungal pathogens.

## Introduction

*Candida albicans* is a commensal fungus that is found in the human oral cavity as well as the gastrointestinal and genitourinary tracts (1). In general, *C. albicans* causes two types of disease in humans. First, mucosal candidiasis of the oral cavity and genitourinary tract are common among both immunocompetent and immunocompromised individuals (2). For example, oral mucocutaneous infections occur in normal newborns while the same disease occurs later in life in patients with reduced T-cell function such as those living with HIV/AIDS. In addition, most women will have at least one episode of vulvovaginal candidiasis in their lifetime. The second type of disease caused by *C. albicans* is invasive disease involving infection of the bloodstream, abdominal organs such as liver or spleen, and the central nervous system (3). The risk of these life-threatening infections is increased in patients with reduced neutrophil number or function; premature infants; and in patients who have undergone extensive abdominal surgery as well as other conditions.

The ability of *C. albicans* to undergo the morphogenetic transition from yeast to hyphae is important for the pathogenesis of both mucosal and invasive infections (4). The formation of hyphae has been correlated with the severity of mucosal disease (5) and with damage to deep organs after dissemination (6). In addition, hyphal formation plays a key role in the establishment of biofilms (7). *C. albicans* biofilms contribute directly to the pathogenesis of mucosal disease and biofilms that form on medical devices such as intravascular catheters contribute indirectly to the pathogenesis of invasive disease (8). Accordingly, understanding the transcriptional networks that regulate hyphal morphogenesis and biofilm formation have been the focus of much research (9, 10).

Nobile et al. initially identified the zinc finger transcription factor Rob1 in a landmark screen for transcription factors required for in vitro biofilm formation and named the gene, **<u>r</u>**egulator **<u>o</u>**f **<u>b</u>**iofilms 1 (9). Rob1 is also required for biofilm formation in vivo in models of intravascular catheter infection and denture infection (9). Recently, we found that loss of *ROB1* function reduces virulence in a model of oropharyngeal candidiasis and decreases filamentation during infection of mammalian tissue (11, 12). As part of our interest in *C. albicans* haploinsufficiency (13), we observed that heterozygous deletion mutants of *ROB1* had distinct filamentation and biofilm phenotypes. This prompted a more detailed analysis of the function of *ROB1* and its two phenotypically distinct alleles.

As described below, we discovered that the reference strain SC5314, and its derivatives are heterozygous at the *ROB1* locus with one allele showing gain-of-function properties relative to the other under some but not all conditions. Curiously, we have been unable to identify the putative gain-of-function allele in any other clinical isolate despite examining 245 strains. This gain-of-function allele likely contributes to the robust in vitro filamentation displayed by strain SC5314 compared to many other strains and may also contribute to the reduced infection of oral tissues that is characteristic of strains with strong in vitro filamentation phenotypes. These results also highlight how differences in the presence of heterozygous, non-synonymous SNPs can contribute significantly to the phenotypic heterogeneity of a diploid eukaryotic pathogen.

## Results

### Rob1 affects hyphal morphogenesis and biofilm formation in an inducing-condition- and strain-dependent manner

The effect of *rob1*ΔΔ mutants on filamentation has been reported in the SC5314-derived SN background in two previous screens of the TF deletion library performed by the Johnson laboratory (9, 14). Homann et al. found that the *rob1*ΔΔ had reduced filamentation on solid agar YPD, YPD+ 10% bovine calf serum (BCS), and Spider medium plates at 37°C (9). We observed the same reduced filamentation phenotype for the library *rob1*ΔΔ strain on Spider medium at 37°C; at 30°C, the mutant showed an altered central wrinkling pattern but peripheral invasion was present (Fig. 1A). Wrinkling in the central portion of the colony is indicative of pseudohyphal growth (15) while the peripheral invasion reflects hyphal growth. At 37°C, the colonies were smooth with no peripheral invasion. On RPMI and RPMI + 10% BCS agar plates, the *rob1*ΔΔ mutant showed strong peripheral filamentation at both temperatures. Within a standard laboratory background, Rob1, therefore, has temperature- and medium-dependent effects on filamentation.

**Figure 1.**
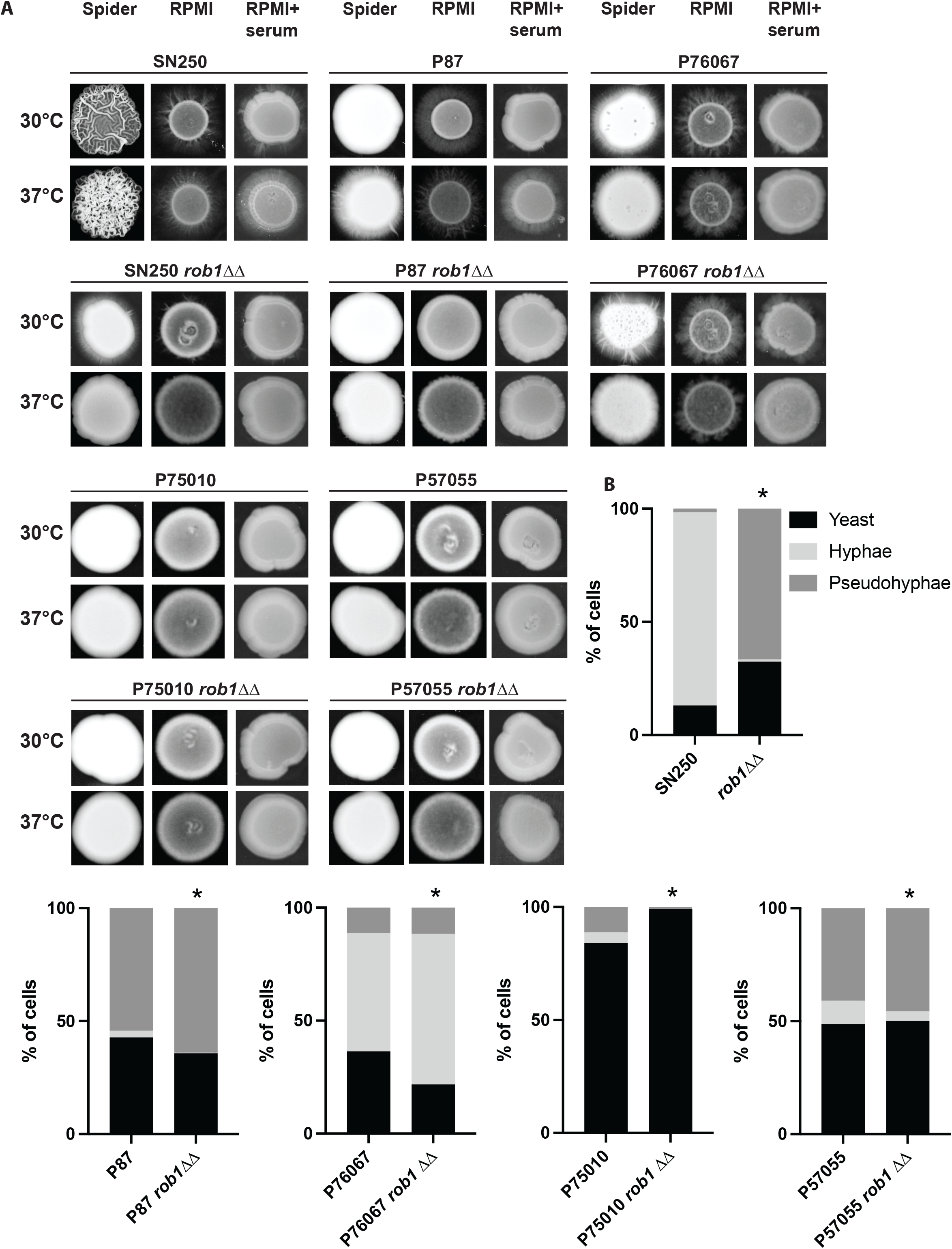
The effect of Rob1 on *C. albicans* filamentation is dependent upon strain background and inducing medium. **A**. Homozygous *ROB1* deletion mutants in the indicated strain backgrounds were spotted on agar plates prepared with Spider medium, RPMI medium, and RPMI supplemented with 10% bovine calf serum (BCS). The strains were incubated at 30°C or 37°C for 3-5 days prior to photographing. Representative images from three independent experiments are shown. **B**. Overnight cultures of the indicated strains were diluted (1:50) into liquid RPMI+10%BCS at 37°C. After 4 hr, the cultures were fixed and the distribution of yeast, pseudohyphae, and hyphae was determined by light microscopy. An asterisk indicates that the parental and *rob1*ΔΔ mutant had statistically significant differences in distribution by Student t test (P < 0.05).

Next, we asked if the effect of Rob1 on filamentation varied with strain background (Fig.1A). To do so, we generated deletion mutants of *ROB1* in four *C. albicans* well-characterized clinical isolates (P75010, P87, P57055, and P76067) for which the effects of other filamentation-related TFs have been studied (16). These four isolates have different filamentation phenotypes: P75010 and P57055 show almost no filamentation while P87 and P76067 filament on RPMI and RPMI + 10% BCS (Fig. 1A). For the low filamenting strains, deletion of *ROB1* has no observable effect except in the case of P75010 on Spider medium at 30°C where the deletion mutant colony shows peripheral invasion. On Spider medium, neither P87 nor P76067 undergo significant peripheral invasion but P76067 shows central wrinkling at 30°C in both WT and the *rob1*ΔΔ mutant but not at 37°C. P76067 has minimal peripheral invasion on Spider medium at 37°C and this invasion is absent in the *rob1*ΔΔ mutant. Deletion of *ROB1* in P87 reduces filamentation at 37°C on both RPMI and RPMI + 10% BCS but not at 30°C. Curiously, P7067 does not filament well on RPMI + 10% BCS but does so on RPMI. The deletion of *ROB1* reduces filamentation of P76067 on RPMI at both 30°C and 37°C. These data indicate that the role of Rob1 during filamentation on agar varies with both the specific induction conditions and the *C. albicans* strain background.

Huang et al. characterized the effect of other key TFs involved in hyphal morphogenesis in these same strains using liquid RPMI+10% BCS conditions (16). Therefore, we examined the effect of the *rob1*ΔΔ mutant under the same conditions. Consistent with the results reported by Huang et al. (16), the 4 clinical isolates as well as the SC5314-derived SN strain showed distinct patterns of filamentation (Fig. 1B). Under these conditions, SN250 and P76067 were the only two strains to form more than 20% true hyphae. P87 and P57055 predominately formed pseudohyphae while >80% of P75010 cells remained as yeast in RPMIS at 37°C for 4 hr (Fig. 1B). Deletion of *ROB1* in SN250 essentially abrogated hyphae formation with pseudohyphae (70%) being the dominant morphotype. In striking contrast, deletion of *ROB1* in P76067 had no effect on hyphae or pseudohyphae formation. Similarly, the *rob1*ΔΔ mutation did not significantly change the morphological distribution of the pseudohyphae-predominate strains P87 or P57055 nor the yeast predominate strain P75010. Huang et al. found that the TFs Efg1, Brg1, and Ume6 had consistent effects across the strains while the role of Bcr1 during filamentation varied between strains (16). Our results indicate that the effect of Rob1 on filamentation is highly condition and strain specific.

In contrast to the condition dependence of Rob1 during *C. albicans* filamentation, it is required for biofilm formation in all conditions reported to date (9, 17). However, the strain dependence of this function had not been explored. For this analysis, we used RPMI medium at 37°C because it supported consistent biofilm formation across all four clinical isolates. We included the SN250 reference strain in each of our experiments to illustrate the relative biofilm formation of the 4 clinical isolates compared to the reference strain under our assay conditions. During the initial adhesion step (Fig. 2), none of the clinical isolates adhered as extensively as SN250 but we also observed no effect of the *rob1*ΔΔ mutant on adhesion in any of the four strain backgrounds. Despite initially adhering to the plastic of the plates, none of the *rob1*ΔΔ mutants were able to develop a biofilm over the next 48 h while the clinical isolates developed reasonable biofilms with P57055 being the poorest biofilm forming strain. Thus, Rob1 regulates biofilm related functions across clinical isolates with both moderate and robust biofilm characteristics.

**Figure 2.**
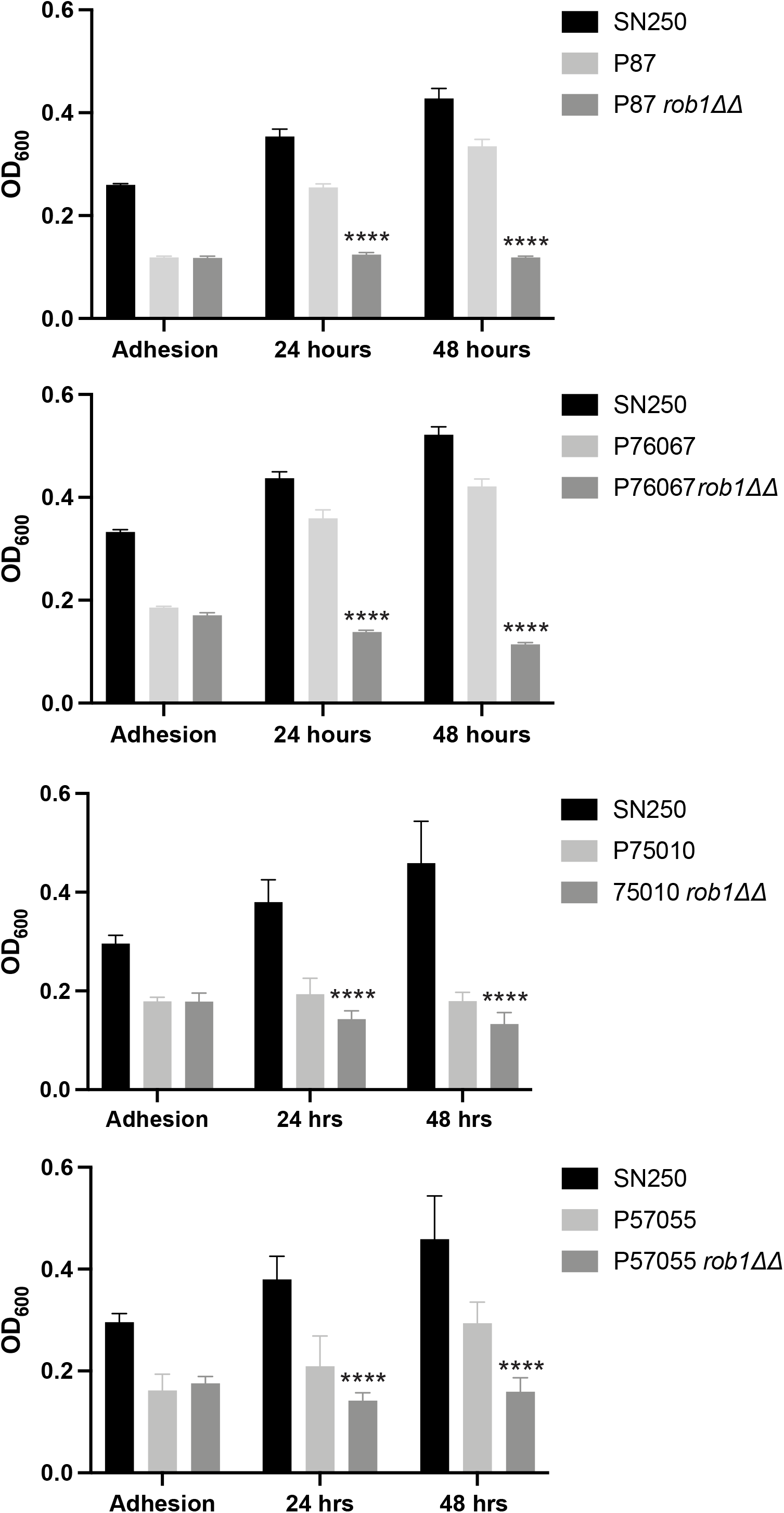
Rob1 is required for biofilm formation in multiple *C. albicans* strain backgrounds. The biofilm formation of the indicated strains was determined using the microtiter plate density assay as described in materials and methods using RPMI medium at 37°C. The bars indicate the OD600 of the biofilm at 90 min (adhesion), 24 hr, and 48hr. The bars indicate mean of biological triplicates performed in technical triplicate. The asterisks indicate that the *rob1*ΔΔ mutant is significantly different from the parental by Student’s t test.

### The set of genes regulated by Rob1 during hypha formation varies with inducing conditions

The effect of Rob1 on gene expression was characterized by Nobile et al. during biofilm formation in Spider medium using microarrays (9) and by Nanostring during kidney and ear infection by Xu et al. (18) and Wakade et al. (12), respectively. However, the role of Rob1 in the regulation of gene expression during in vitro hyphal induction conditions has not been reported. Using an RNA sequencing approach, we compared gene expression of *rob1*ΔΔ mutants to WT after 4 hr of hyphal induction using RPMI and Spider medium at 37°C. A total of 211 genes were differentially expressed (±log_2_ 1; adjusted p value <0.05) in the *rob1*ΔΔ mutant in Spider medium and 263 genes were differentially expressed in RPMI medium relative to SN250 (Fig. 3A&B). Strikingly, only 10 genes were down regulated in the *rob1*ΔΔ mutant in both media and there were no genes upregulated in both media (Fig. 3C&D). The set of Rob1-dependent genes downregulated under both conditions contained well-known hypha specific genes (*HWP1*, *HYR1*, *ECE1*, and *SAP6*) as well as the recently described TF involved in biofilm formation, *WOR3* (19).

**Figure 3.**
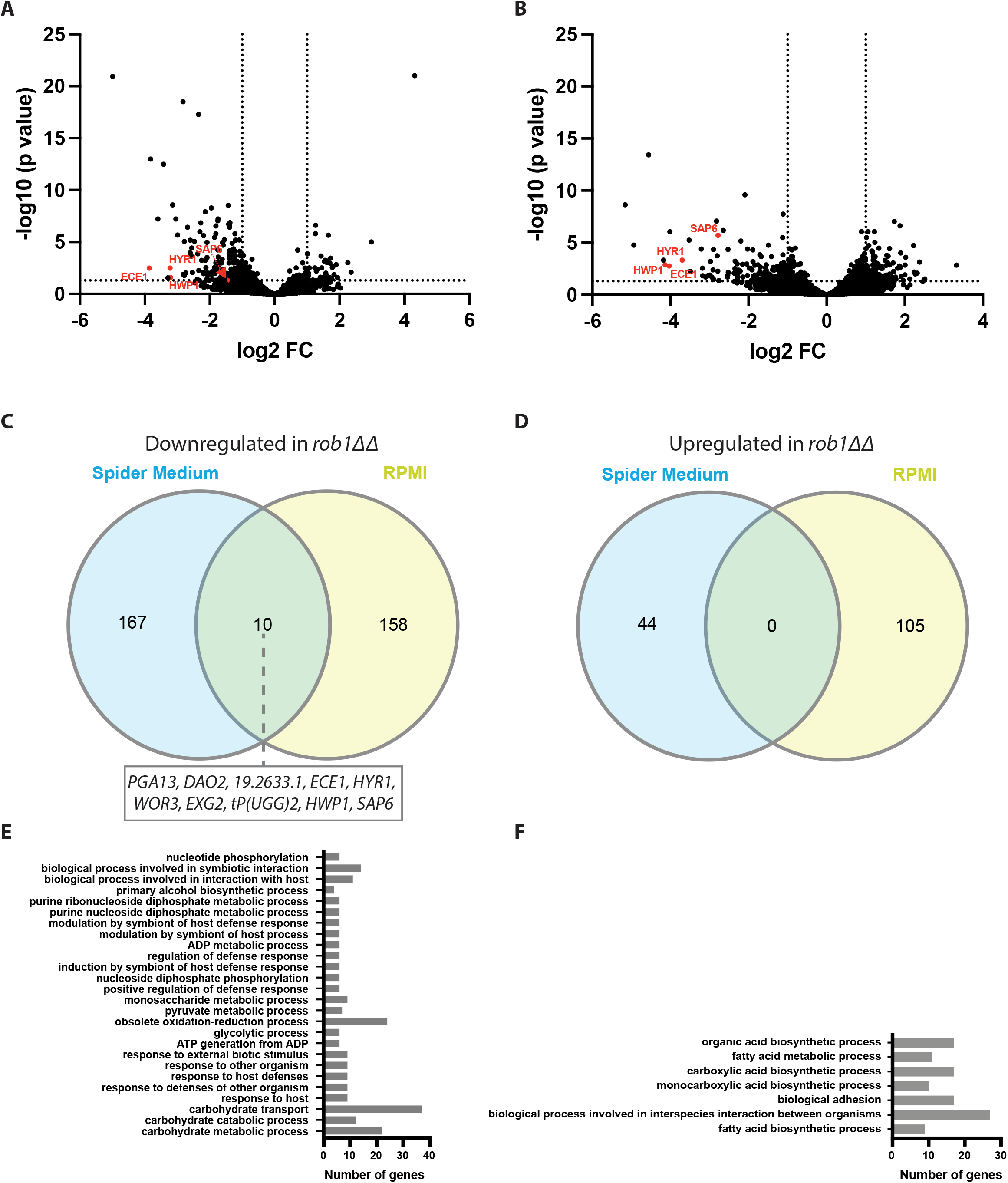
The Rob1 regulon during filamentation is dependent upon the inducing conditions. Volcano plots for differentially expressed genes (± 1 log_2_, adjusted P value < 0.05) between the SN250-derived *rob1*ΔΔ mutant and SN250 in Spider medium (**A**) and RPMI (**B**) at 37°C for 4hr. Genes indicated in red are hyphae-associated genes. Venn diagrams comparing the genes downregulated (**C**) or upregulated (**D**) in the two media. GO terms for genes differentially expressed in the *rob1*ΔΔ mutant in Spider medium (**E**) and RPMI medium (**F**).

GO term analysis of the set of genes downregulated in the *rob1*ΔΔ mutant in Spider medium identified carbohydrate-related genes including transport, glycolysis and pyruvate metabolism as the top classes of genes (Fig. 3E). In contrast, genes involved in fatty acid and carboxylic biosynthesis, ergosterol biosynthesis, adhesion and biofilm formation are enriched in the set of genes downregulated in RPMI (Fig. 3F). In Spider medium, amino acid and nucleotide transport genes were upregulated in the *rob1*ΔΔ mutant while in RPMI autophagy genes were induced. These data suggest that the specific sets of genes dependent on Rob1 for full expression during hyphal induction vary substantially depending on the nature of the medium used for induction.

The two categories of genes enriched in the conditions represent two different pathways of carbon metabolism: carbohydrate metabolism in Spider medium and fatty acid/lipid metabolism in RPMI. Spider medium is comprised of beef extract nutrient broth and mannitol; as such, it lacks glucose; RPMI contains glucose but does not have added lipid or fatty acids. One possible explanation for these observations is that Rob1 responds to the carbon sources available to cell during hyphae morphogenesis. However, the *rob1*ΔΔ mutant does not have reduced growth in these media and, therefore, it does not seem to be a generalized regulator of carbon metabolism. Because the *rob1*ΔΔ mutant in the SN background filaments in RPMI medium but not on Spider medium (Fig. 1A), it seems that the reduced expression of fatty acid metabolism genes in *rob1*ΔΔ may be less important to the process of filamentation than reduced carbohydrate metabolism.

### SC5314-derived strains are heterozygous at the *ROB1* locus and the two alleles have distinct filamentation phenotypes

We previously constructed a library of TF heterozygotes for use in genetic interaction studies (13). As part of this work, we observed that *ROB1* heterozygous deletion mutants showed distinct filamentation phenotypes on Spider medium at 37°C. The phenotypes of six independent *rob1*Δ/*ROB1* heterozygotes constructed in the SN/SC5314 background are shown in Fig. 4A along with the parental strain and the *rob1*ΔΔ homozygote. Three heterozygotes show wild type or even increased levels of filamentation after 7 days on Spider medium at 37°C. The other three heterozygotes show smooth colonies with little or no peripheral invasion and are closer to the null mutant. As indicated in the Candida Genome Database, *ROB1* in SC5314 is heterozygous with a non-synonymous C/T polymorphism at position 2902 resulting in the A allele encoding a serine and the B allele encoding a proline (Fig. 4B). The results of Sanger sequencing of this region of *ROB1* for the six isolates is shown in Fig. 4A and the genotypes are as indicated in Fig. 4B. The filamentous isolates *(rob1-2*, *rob1-3*, *rob1-5, rob1-6*) contained an S at 946 (*ROB1^946S^*/*rob1^946P^*Δ) while the isolates with reduced filamentation *(rob1-1*, *rob1-4*) contained P at 946 (*rob1^946S^Δ*/*ROB1^946P^*). For the remainder of our studies, we used strains *rob1-1* to characterize the *rob1^946S^Δ*/*ROB1^946P^* genotype and *rob1-5* for the *ROB1^946S^*/*rob1^946P^*Δ genotype.

**Figure 4.**
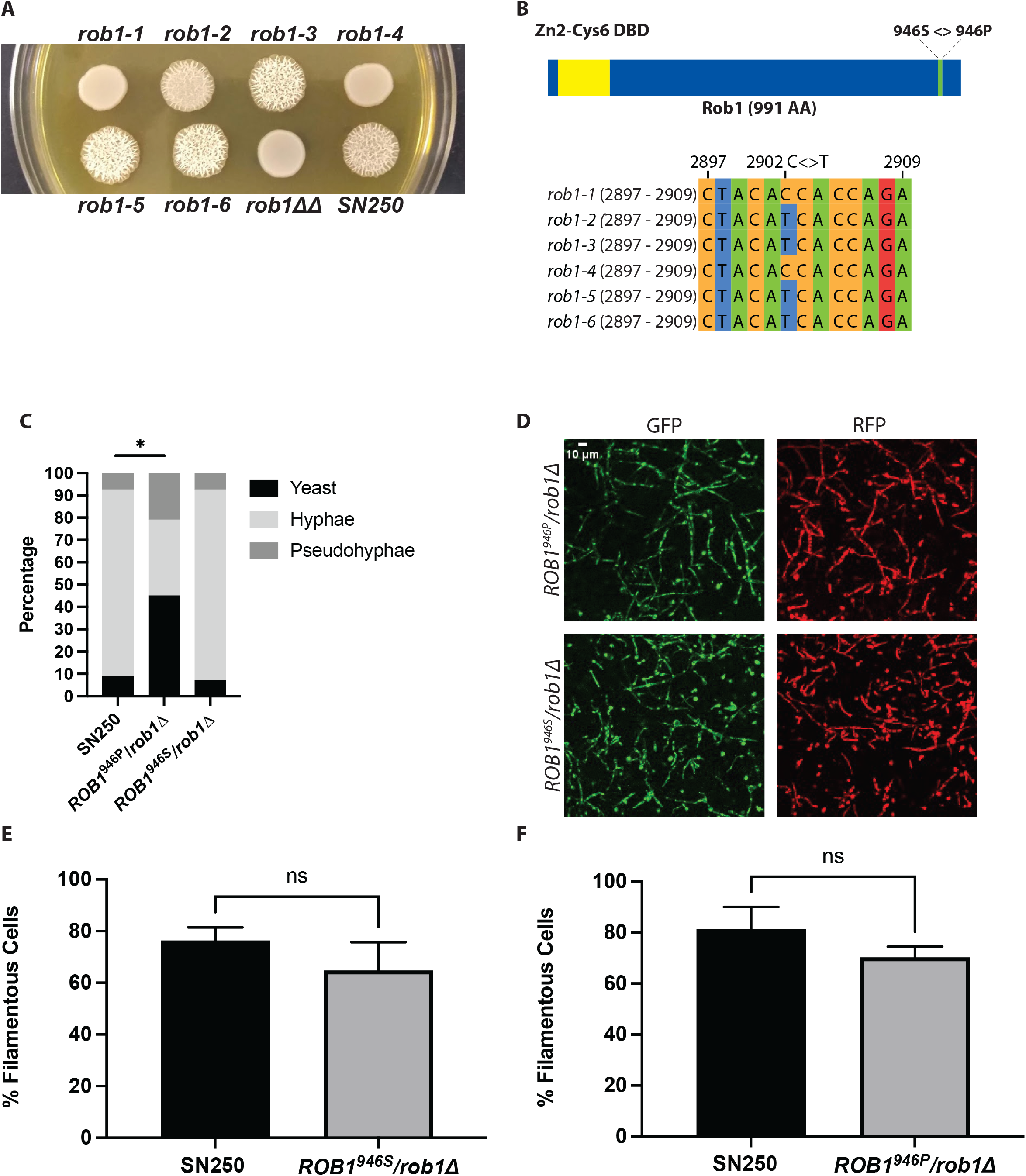
*ROB1* in the SC5314 background is heterozygous and the two alleles have different filamentation phenotypes. **A**. Six independent isolates of heterozygous *ROB1* strains derived in the SN background were plated on solid Spider medium and incubated for 4 days at 37°C. **B**. Diagrams of the Rob1 ORF indicating the position of the SNP and the Sanger sequences of the six heterozygous strains shown in (A). **C**. The distribution of cell morphologies after 4 hr incubation in RPMI+ 1%BCS. The asterisk indicates that the indicated strain differs from SN250 in a statistically significant manner by Student’s t test (P < 0.05). **D**. Micrographs of of mouse ears inoculated with a 1:1 mixture of SN250 (GFP labelled) and the indicated *rob1* mutant (RFP) and imaged by confocal microscopy 24 hr post infection. Quantitation of the percentage of filamentous cells for the two strains (**E**&**F**). For details of imaging and quantitation see materials and methods. NS indicates the differences were not significant by Student’s t test (P >0.05).

To determine if the filamentation phenotype of the heterozygous *ROB1* mutants varied with inducing conditions, the isolates *rob1-1* and *rob1-5* were plated on RPMI and RPMI+ 10% BCS medium and incubated at both 30°C and 37°C. Under these conditions, there was no difference between the filamentation of either heterozygous mutant and the reference strain at either temperature (Supplementary Fig. 1A); on Spider medium at 30°C the *rob1-1* isolate showed reduced wrinkling relative to the reference strain and the *rob1-5* isolate. In liquid inducing conditions with RPMI+ 10% BCS at 37°C, we observed no difference in the distribution of yeast/hyphae/pseudohyphae between the two *ROB1* heterozygotes and the reference strain (Supplementary Fig.1B). However, when the amount of BCS was reduced from 10% to 1%, the *rob1-1* mutant generated less hyphae than the reference strain and *rob1-5* and a corresponding increased proportion of pseudohyphae and yeast (Fig. 4C). These observations indicate that the *ROB1^946S^*/*rob1^946P^*Δ strain appears carry a phenotypic gain-of-function allele relative to the *rob1^946S^Δ*/*ROB1^946P^*strain under specific filamentation conditions.

Finally, we compared the filamentation of the two *ROB1* alleles in vivo using an intravital imaging assay based on infection of the subcutaneous tissue of the mouse ear (20). We have previously shown that the *rob1*ΔΔ mutant has dramatically reduced filamentation in this model (12). In this experiment, the *ROB1^946S^*/*rob1^946P^*Δ and *rob1^946S^Δ*/*ROB1^946P^* strains were labelled with iRFP and each injected into the ear tissue as a 1:1 mixture with SN250 that had been labelled with GFP. Twenty-four hours post infection, the extent of filamentation (% filamentous cells) and the length of the filaments for the heterozygous *ROB1* strains were compared to the WT strains. Both heterozygous strains formed filaments to a similar extent as WT in the ear (Fig. 4D-F). The lengths of each filament were also similar (Supplementary Fig. 1C). This indicates that the filament-inducing stimuli in mammalian tissue are sufficient to trigger filamentation in the *rob1* heterozygotes regardless of the allele that is present.

### The *ROB1^946P^* /*rob1*Δ strain has reduced biofilm formation in vivo and in vitro

Next, we examined the effect of the two *ROB1* alleles on biofilm formation under three conditions, Spider medium, RPMI+1% BCS, and RPMI+10% BCS at 37°C. The *rob1^946S^Δ*/*ROB1^946P^*strain showed reduced biofilm relative to both wild type and the *ROB1^946S^*/*rob1^946P^*Δ strain under all three conditions (Fig. 5A) during adhesion and at 24 hr and 48 hr. The *rob1^946S^Δ*/*ROB1^946P^* strain phenocopied the *rob1*ΔΔ under all three conditions, indicating it is significantly attenuated in biofilm formation in vitro. We also examined the structure of the biofilms formed in Spider medium and found that the biofilm formed by the *rob1^946S^Δ*/*ROB1^946P^* strain was less dense than either WT or the *ROB1^946S^*/*rob1^946P^*Δ strain but that there was no difference in the presence of hyphae within the biofilm structure (Fig. 5B).

**Figure 5.**
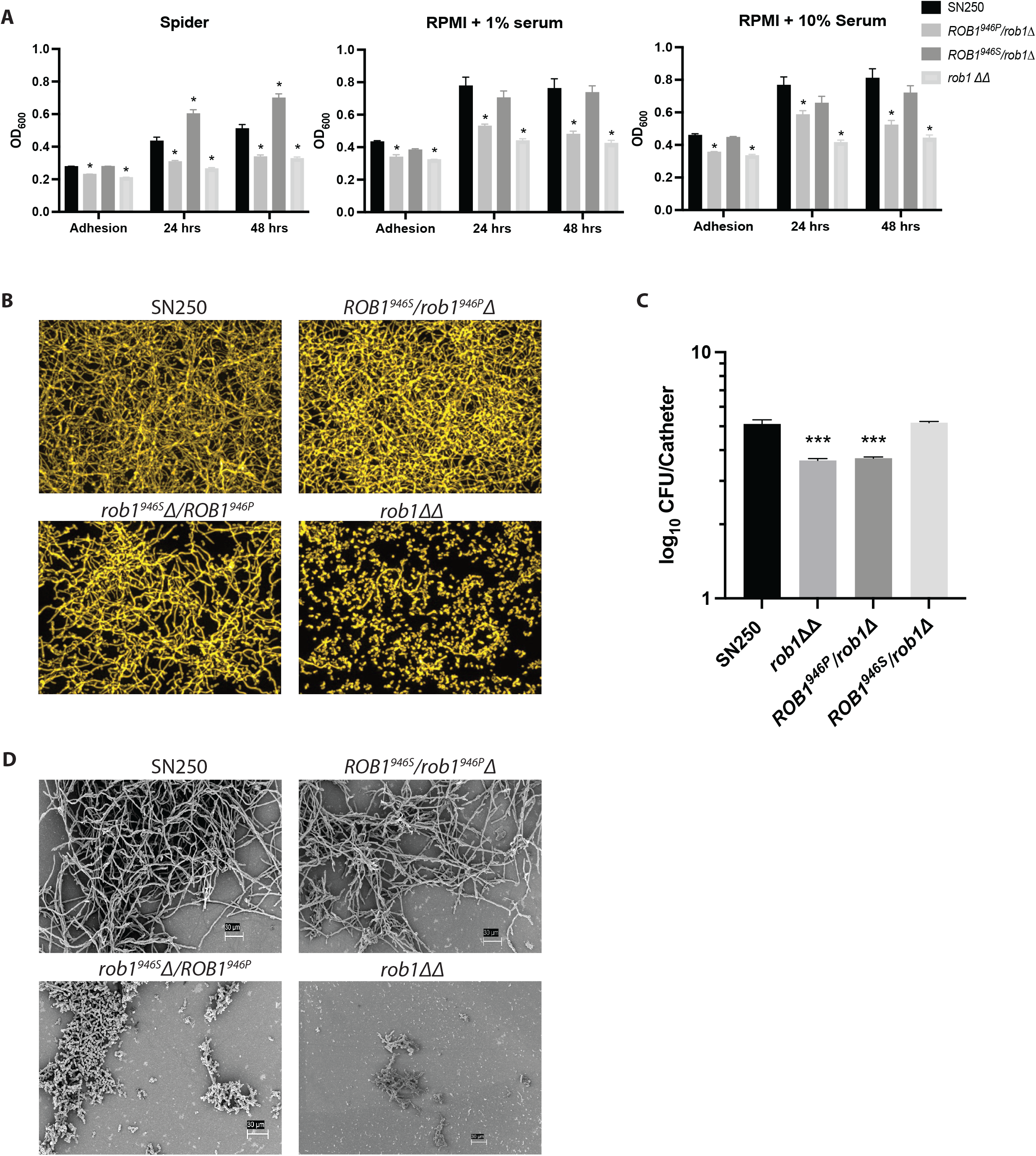
The *ROB1* alleles have distinct biofilm formation phenotypes in vitro and in vivo. **A**. The biofilm formation of *ROB1* heterozygotes was compared to SN250 and the *rob1*ΔΔ mutant in Spider medium, RPMI+1% serum, and RPMI+10% serum at the indicated time points. The asterisks indicate statistically significant changes from SN250 using Student’s t test corrected for multiple comparisons (adjusted P <0.05). **B**. The biofilms were imaged using a microtiter plate imaging system as described in materials and methods. The apical views are shown and are representative of two replicates. **C**. The fungal burden of intravascular catheters infected with the indicated strains 24 hr post infection. The bars indicate mean fungal burden from catheters placed in three rats and the error bars indicate standard deviation. The asterisk indicates statistically significant differences from WT by ANOVA followed by Dunnett’s correction for multiple comparisons (adjusted P value <0.05). **D**. Scanning electron microscopy of biofilms formed by the indicated strains in the vascular catheters 24 hr post infection.

Nobile et al. showed previously that the *rob1*ΔΔ mutant has reduced biofilm formation in a rat intravascular catheter infection model (9). To determine if the phenotypes observed in vitro were also present in vivo, catheters implanted in jugular veins of rats were infected with SN250*, ROB1^946S^*/*rob1^946P^*Δ, *rob1^946S^Δ*/*ROB1^946P^*, and *rob1*ΔΔ strains. The catheters were removed 24 hr post infection and the fungal burden was determined as previously described. Consistent with previous results, catheters infected with the *rob1*ΔΔ mutant had reduced fungal burden (∼1.5 log_10_ CFU/catheter) relative to SN250 (Fig. 5C). The fungal burden of the catheters infected with the *ROB1^946P^*/*rob1*Δ strain was nearly identical to the *rob1*ΔΔ mutant while catheters infected with the *ROB1^946S^*/*rob1*Δ strain was comparable to SN250. The infected catheters were also characterized using scanning electron microscopy. The *ROB1^946S^*/*rob1*Δ strain formed hyphal structures with a structure similar to the SN250 reference strain (Fig. 5D). The *rob1^946S^Δ*/*ROB1^946P^* strain, on the other hand, forms biofilm structures that consist mainly of yeast with some cells that appear to be pseudohyphae. It is interesting to note that in Spider medium, the *rob1^946S^Δ*/*ROB1^946P^* strain forms less dense biofilms that are structurally similar to WT (Fig. 5B), while in vivo (Fig. 5D), the biofilm shows a dramatic reduction in filamentous forms.

### The *ROB1^946S^* allele appears to be rare and isolated to the SC5314 strain among 224 sequenced isolates

Next, we were interested to determine the prevalence of the *ROB1^946P^*allele. To identify other strains with this SNP, we compiled genomes from 216 *C. albicans* strains from the literature (21, 22, 23, 24) and an additional 8 strains collected from premature infants that our group had sequenced (25). SNP calling was performed as described in the materials and methods. Although we do not have uniform sampling from all clades, we have been unable to identify another *C. albicans* strain that contains the *ROB1^946S^* allele as either a heterozygote or a homozygote (Table 1). In other words, all sequenced strains that we have analyzed to date are homozygous for the *ROB1^946P^* allele which is the phenotypically less active allele of *ROB1*. Our largest set of sequenced genomes comes from Clade 1, which includes SC5314. Thus, the *ROB1^946S^* allele is rare among relatively closely related strains. In previously reported systematic studies of filamentation and biofilm formation across a large set of clinical isolates (22, 26), SC5314 is one of the most robust in terms of these two phenotypes under many conditions. It is therefore tempting to speculate that the *ROB1^946S^* allele may contribute to this feature of the strain.

**Table 1.** Analysis of *ROB1* sequence in 253 *C. albicans* isolates. **Summary of *ROB1* position 2902 genotypes**

| Clade (MLST) | Number of strains analyzed | 2902 C -> T SNP present | Strains with the SNP |
| --- | --- | --- | --- |
| 1 | 74 | Yes | SC5314 |
| 2 | 16 | No |  |
| 3 | 18 | No |  |
| 4 | 31 | No |  |
| 5 | 1 | No |  |
| 6 | 1 | No |  |
| 7 | 1 | No |  |
| 8 | 4 | No |  |
| 9 | 4 | No |  |
| 10 | 4 | No |  |
| 11 | 11 | No |  |
| 12 | 6 | No |  |
| 13 | 23 | No |  |
| 14 | 0 | N/A |  |
| 15 | 0 | N/A |  |
| 16 | 2 | No |  |
| 17 | 0 | N/A |  |
| 18 | 4 | No |  |
| NC | 24 | No |  |
| Total strains analyzed | 224 | Total strains with the 2902 C -> T SNP | 1 |

The position of the non-synonymous SNP is in the C-terminal portion of the protein. Rob1 is a zinc finger transcription factor and its likely DNA binding domain is predicted to be in the C-terminal region of the protein (27). Gain-of-function mutations in zinc finger transcription factors frequently are located in the C-terminus of the protein. For example, fluconazole resistance is associated with such mutations in the zinc finger transcription factors Tac1 and Mrr1 (28); these mutations lead to increased expression of multi-drug efflux pumps such as *CDR1* which mediate fluconazole resistance. Our initial phenotypic data suggest that *ROB1^946S^* may represent a gain-of-function allele relative to *ROB1^946P^* which is the predominant allele in sequenced *C. albicans* isolates.

Although there are a variety of mechanisms by which an allele can display phenotypes of a gain-of-function allele, the simplest and best characterized for a transcription factor is that the changes in amino acids alter the activity of the factor. As discussed above, the Rob1 946 position is in the C-terminus which is frequently the activation domain of zinc finger transcription factors. If that were the case, then we would expect that one allele would activate the expression of genes regulated by Rob1 more than the other. We focused our analysis on three canonical hypha-associated genes (*ALS3, ECE1,* and *HWP1*) differentially expressed in the *rob1*ΔΔ mutant. We also examined the expression of *ROB1* to see if the alleles may auto-regulate the gene differently. We examined the expression of these four genes during hyphal induction by RPMI with 1% BCS because the mutants have distinct filamentation phenotypes under these conditions (Fig. 6).

**Figure 6.**
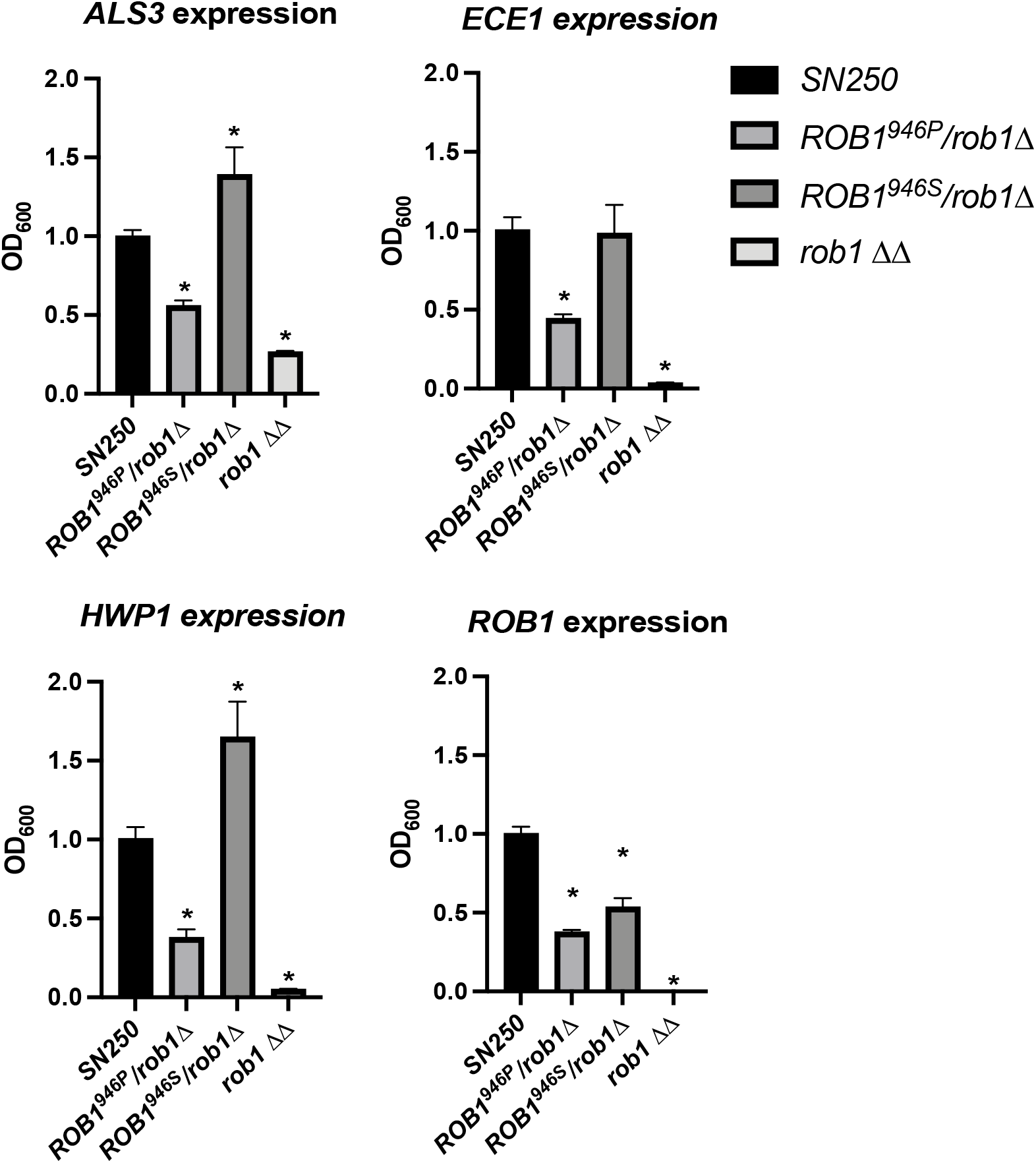
The *ROB1* alleles have distinct effects on the expression of canonical hyphae-associated genes during filamentation. The indicated strains were incubated in RPMI+1%BCS for 4 hr and RNA isolated as described in the materials and methods. The expression of *ALS3*, *ECE1*, *HWP1*, and *ROB1* were determined by quantitative RT-PCR using the ΔΔCT method. The bars indicate mean of two independent experiments preformed in triplicate. Asterisks indicate statistically significant differences between the indicated strain and SN250 by ANOVA followed by Dunnett’s correction for multiple comparisons (P <0.05).

We first compared the heterozygous mutants to WT and the *rob1*ΔΔ mutant under these conditions. *ROB1* expression was reduced by ∼2-fold in both heterozygous mutants relative to WT (Fig. 6). The similar expression of *ROB1* in the two heterozygous strains indicates that phenotypic differences are not due to differential autoregulation but more likely due to differences in the function of the resulting protein. For *ALS3*, *ECE1*, and *HWP1*, a consistent pattern of relative gene expression emerged. The three genes were expressed in the *ROB1^946S^*/*rob1^946P^*Δ mutant at a higher level compared to the *rob1^946S^*Δ/*ROB1^946P^*mutant. The expression of the genes in the WT strain was intermediate between the two heterozygous mutants (*HWP1* and *ALS3*) or was comparable to the *ROB1^946S^*/*rob1^946P^*Δ mutant (*ECE1*). Although the changes in gene expression between the two alleles are not dramatic (2-3 fold), they correlate with the distinct filamentation phenotypes shown by the strains and support the conclusion that the rare *ROB1^946S^*allele found in SC5314 represents a gain-of-function allele relative to the more prevalent *ROB1^946P^* allele.

### Introduction of the *ROB1^946S^* allele increases filamentation in both strong and weak filamenting strain backgrounds

We were interested to test the hypothesis that *ROB1^946S^* is a gain-of-function allele by an allele swap of *ROB1^946P^*with *ROB1^946S^* in a strain homozygous for the *ROB1^946P^.* If this hypothesis is correct than we expected to observe an increase the ability of the allele swapped strain to undergo filamentation relative to the parental strain. We replaced one allele of a poorly filamenting, *ROB1^946P^*homozygous clinical strain (RO39) with *ROB1^946S^* carrying a *NAT1* marker. We also constructed a homozygous *ROB1^946P^* strain with a *NAT1* maker in the same chromosomal position as the allele swap mutant for a control (Supplementary Fig.2 with construct). Neither the parental strain nor its Nat+ derivative filamented well on either solid RPMI or Spider medium (Fig. 7A). In contrast, the *ROB1^946P^/ ROB1^946S^* strain showed both wrinkling and peripheral invasion on Spider medium and peripheral invasion on RPMI medium. In liquid RPMI + 10% BCS medium (Fig. 7B), the homozygous *ROB1^946P^/ ROB1^946P^* mutant formed 31% filaments after 4 hr at 37°C while the heterozygous derivative *ROB1^946P^/ ROB1^946S^* formed 51% filaments. RO39 forms very poor biofilms but introduction of the *ROB1^946S^*allele did not affect biofilm formation under any of the conditions tested (Supplementary Figure 2A). These data support the conclusion that the SC5314-derived *ROB1^946S^* allele is a gain-of-function allele relative to the *ROB1^946P^*allele but its phenotypic effects appear to be condition and strain dependent.

**Figure 7.**
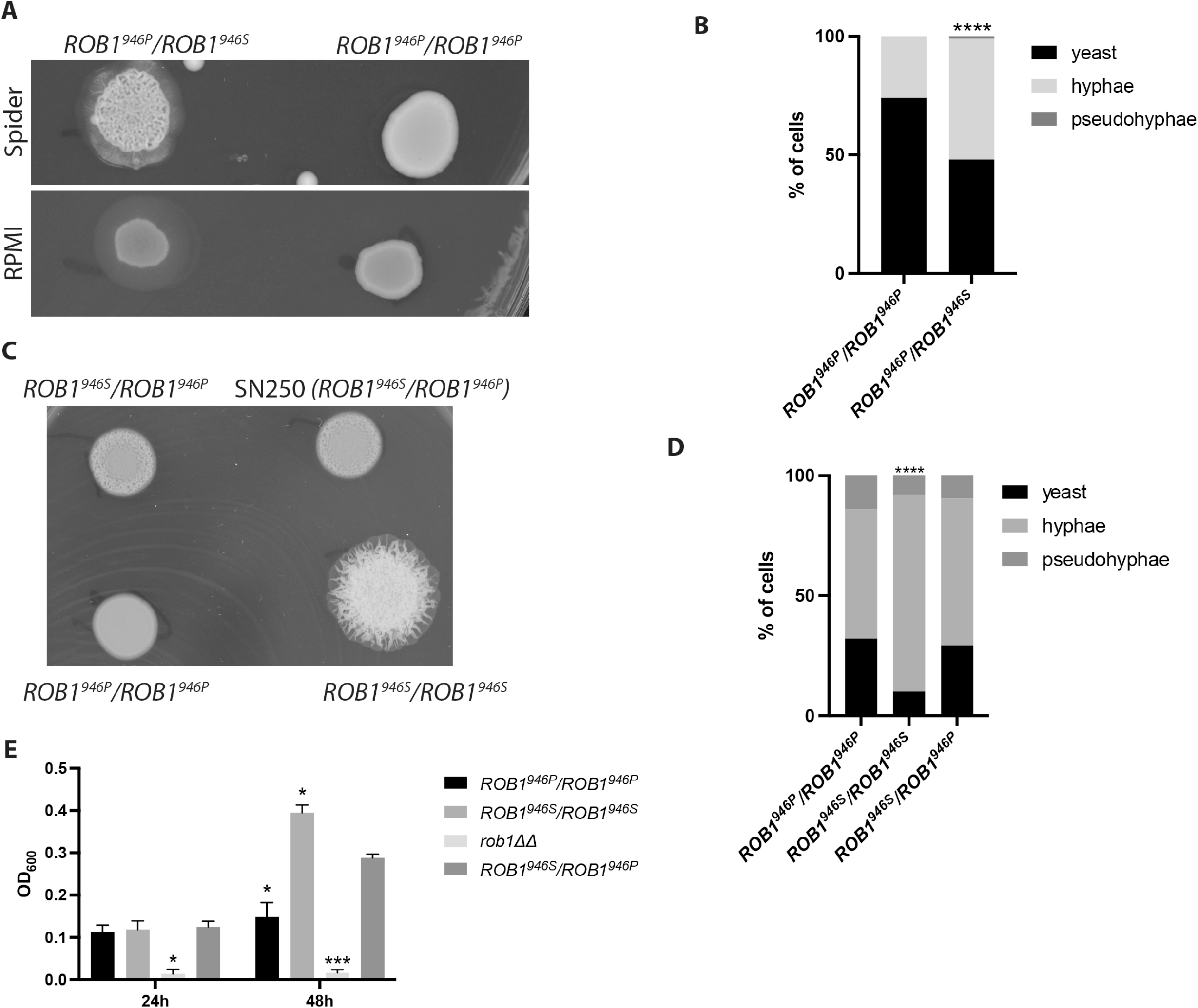
The *ROB1^946S^*allele appears to be a gain-of-function allele during in vitro filamentation and biofilm formation. **A**. A poorly filamenting clinical isolate of *C. albicans* that is homozygous for the *ROB1^946P^*allele was converted to an SC5314-like *ROB1* heterozygote by knock-in of the *ROB1^946S^* allele. The resulting strain shows increased filamentation relative to a strain with knock-in of the exogenous allele on solid Spider medium and RPMI (**A**) and after induction in liquid RPMI+1%BCS for 4 hr at 37°C (**B**). The asterisk indicates that the indicated strain differs from SN250 in a statistically significant manner by Student’s t test (P < 0.05). C. SN250 was transformed with knock-in constructs to generate either *ROB1* homozygotes or a heterozygote containing NAT markers in the 3’ untranslated region. These filamentation of these strains were compared to the unmarked SN250 heterozygote on Spider medium at 37°C (**C**) and in liquid RPMI+10% BCS (**D**). The asterisk indicates that the indicated strain differs from SN250 in a statistically significant manner by Student’s t test corrected for multiple comparisons (P < 0.05). **E**. The biofilm properties of the SN250 knock in strains were compared in RPMI+1% BCS at 37°C at 24 and 48 hr. Asterisks indicate statistically significant differences between the indicated strain and SN250 by ANOVA followed by Dunnett’s correction for multiple comparisons (P <0.05).

To further test this hypothesis, we constructed homozygous *ROB1^946P^/ ROB1^946P^* and *ROB1^946S^/ ROB1^946S^* strains in the SC5314-derived SN background using a CRISPR-Cas9-based approach. As with the single allele swap strains, the parental heterozygous strain with a *NAT1* marker in the 5’ region was also constructed. As shown in Fig. 7C, the homozygous *ROB1^946S^/ ROB1^946S^* strain shows a striking increase in filamentation on Spider medium at a time point when the heterozygous strain shows only the beginning of central wrinkling and the homozygous *ROB1^946P^/ ROB1^946P^*strain shows a smooth colony. Once again, this increase in filamentation is condition dependent because there is no difference in the colonies of the different *ROB1* strains on solid RPMI medium. In RPMI+1% BCS at 37°C (Fig. 7D), the *ROB1^946S^/ ROB1^946S^* strain forms slightly more hyphae after 4hr compared to either the *ROB1^946P^/ ROB1^946P^*or *ROB1^946P^/ ROB1^946S^* strains. The homozygous *ROB1^946S^/ ROB1^946S^* strain in Spider medium (Fig. 7E) shows increased biofilm density in Spider medium with the *ROB1^946P^/ROB1^946S^* heterozygote intermediate between the two homozygous strains.

### The more prevalent *ROB1^946P^* allele promotes a commensal phenotype in oral pharyngeal infection while the SC5314-derived *ROB1*^946S^ allele promotes invasion

To our knowledge, the virulence of the *rob1*ΔΔ mutant has not been examined in the standard mouse model of disseminated candidiasis. Xu et al. examined the effect of the *rob1*ΔΔ mutant on gene transcription in this model and found that kidneys infected with the *rob1*ΔΔ mutant had reduced overall fungal RNA (18), strongly indicating that strain had attenuated virulence. We compared the virulence of the *rob1*ΔΔ mutant to WT and the heterozygous *ROB1^946S^*/*rob1^946P^*Δ and *rob1^946S^*Δ/*ROB1^946P^* strains (Fig. 8A) in the same model of disseminated candidiasis. The *rob1*ΔΔ mutant was significantly attenuated while both heterozygous strains were similar to WT. Although filamentation does not always correlate with virulence (15), the *rob1*ΔΔ mutant showed reduced filamentation in vivo (12) while the heterozygous mutants filamented to the same extent as the WT strain (Fig. 4D-F). The virulence of SN250 derived strains that are homozygous for the ROB1 alleles had the same virulence as the heterozygous strain (Fig. 8B), further indicating that the two alleles have no effect on disseminated candidiasis.

**Figure 8.**
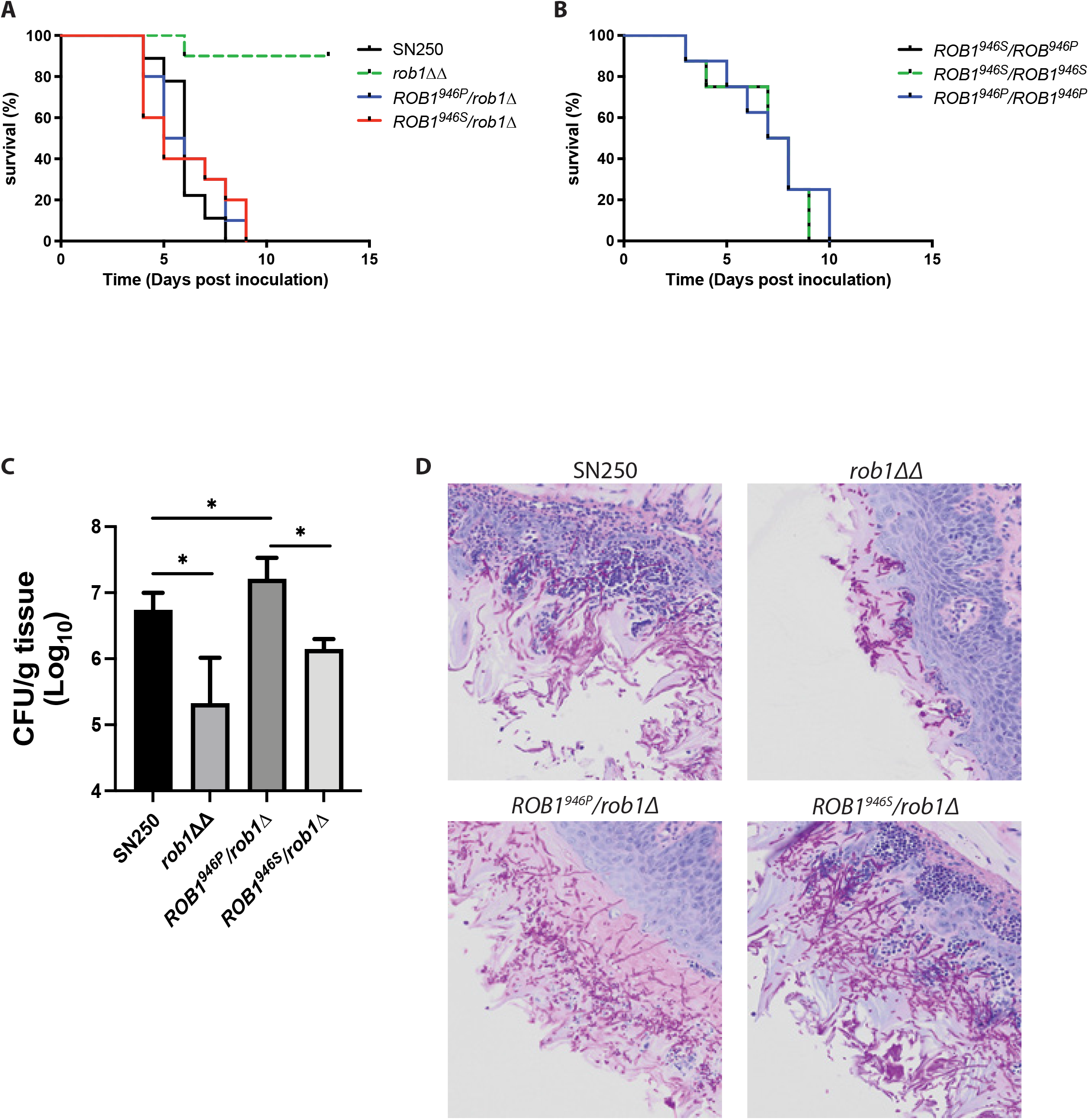
The *ROB1^946P^*allele promotes oral colonization while the *ROB1^946S^* allele promotes invasive infection. **A**&**B**. Survival curves for CD-1 mice (n = 10/strain) infected with the indicated strains by tail-vein infection and monitored to moribundity. The *rob1*ΔΔ mutant (A) was the only strain for which a statistically significant survival time was observed (Kaplan Meier, Mantel Log-Rank, P<0.05). **C**. The oral fungal burden of tongues harvested from cortisone-treated CD-1 mice (5/strain) infected with the indicated strains five days post-infection. The bars are mean with standard deviation indicated by the error bars. The asterisks indicate statistically significant differences between strains denoted by the horizontal lines as determined by ANOVA with Dunnett’s test of multiple comparisons. **D.** Histological analysis of tongues from infections described in (C). The fields are representative of multiple fields evaluated.

Oropharyngeal candidiasis is an infection which shares many pathobiological features with biofilm formation (2, 8). We have previously shown that the *rob1*ΔΔ has reduced virulence in a mouse model of oropharyngeal candidiasis (11). Therefore, we tested the virulence of the heterozygous *ROB1^946S^*/*rob1^946P^*Δ and *rob1^946S^*Δ/*ROB1^946P^* strains in comparison to both the *rob1*ΔΔ mutant and wild type (Fig. 8C). Surprisingly, the fungal burden of tissue from animals infected with the *rob1^946S^*Δ/*ROB1^946P^*mutant was 1 log_10_ CFU/g tongue tissue higher than the animals infected with the *ROB1^946S^*/*rob1^946P^*Δ mutant. The fungal burden of animals infected with the *ROB1^946S^*/*rob1^946P^*Δ mutant was intermediate between the *rob1*ΔΔ mutant and the *rob1^946S^*Δ/*ROB1^946P^*mutant. The differences between the fungal burden established by the *rob1*ΔΔ and *rob1^946S^*Δ/*ROB1^946P^* mutants and WT was statistically significant (p <0.05 by ANOVA and post hoc Student’s t test with Bonferroni correction for multiple comparisons). The difference in fungal burden between animals infected with the *ROB1^946S^*/*rob1^946P^*Δ mutant and WT was not statistically significant. This is the opposite pattern compared to the effects of the different alleles in the catheter model.

To further explore possible explanations for the distinct phenotypes of the *ROB1* alleles during oropharyngeal infection, we examined the histology of the tongues infected with the different *ROB1* heterozygotes (Fig. 8D). Tissue infected with the WT strain shows extensive filamentation of the fungus with invasion into the subepithelial compartment and recruitment of inflammatory cells. A similar phenotype is exhibited by the *ROB1^946S^*/*rob1^946P^*Δ mutant. In stark contrast, tissue infected with the *rob1^946S^*Δ/*ROB1^946P^*mutant is devoid of fungus in the sub-epithelium while the epithelium displays a robust infection of filamentous fungal cells. Furthermore, there is essentially no signs of inflammatory cell recruitment to the sub-epithelial tissue. This non-inflammatory, epithelium-localized infection is also seen in tissue inoculated with the *rob1*ΔΔ mutant. These data suggest that the increased fungal burden observed for the *rob1^946S^*Δ/*ROB1^946P^*mutant relative to the *ROB1^946S^*/*rob1^946P^*Δ mutant is because it establishes a non-inflammatory colonization more consistent with a commensal state (5) while the *ROB1^946S^*/*rob1^946P^*Δ mutant causes a more invasive infection that leads to tissue damage and clearance of the fungus by the inflammatory response. Since the *rob1*ΔΔ mutant is significantly less able to establish infection or colonization, Rob1 is critical for OPC but whether that leads to a commensal or invasive infection depends, in part, on the specific *ROB1* allele.

## Discussion

Here, we characterized the role of Rob1 in two virulence traits, filamentation and biofilm formation, and in three in vivo models of infection: disseminated disease, OPC, and vascular catheter. In vitro, the zinc finger transcription factor Rob1 indicates in *C. albicans* pathobiology that is highly dependent upon the culture media in vitro. An excellent example of this distinction is that the *rob1*ΔΔ mutant forms filamentous colonies on RPMI medium agar plates that show clear invasion, while incubation in the same medium under liquid conditions almost no hyphae are formed and pseudohyphae predominate. Consistent with this environmentally contingent phenotype is our observation that Rob1 regulates distinct sets of genes during in vitro hyphal morphogenesis in two commonly used induction media: Spider medium and the tissue culture medium RPMI (Fig. 3C&D). The overlap in regulated genes is a small fraction of the total number of differentially expressed under the two conditions.

Furthermore, Nobile et al found that 2150 genes are differentially expressed in the *rob1*ΔΔ mutant during biofilm formation which is 10-fold higher than the number of genes we found to be differentially expressed under hyphae induction (9). Based on CHiP-ChiP analysis only 2% of the differentially expressed genes in the *rob1*ΔΔ mutant are bound by Rob1. Thus, Rob1 appears to have indirect effects on gene transcription of a large set of genes during biofilm formation and the indirect nature of its function is one possible explanation for the differences in regulated genes between the two inducing conditions. Until the transcriptional profiles of more TFs involved in filamentation are directly compared under different conditions, it is not possible to know if such a striking change in differentially expressed genes is general phenomenon or specific to Rob1. We have recently compared the transcriptional profiles of *EFG1*, *ROB1* and *BRG1* mutants in RPMI+10%BCS and during infection of ear tissue using Nanostring probe set of 185 environmentally responsive genes (12). Although each gene regulated distinct genes sets in the two conditions, the overlap was much more extensive than observed for the *rob1*ΔΔ mutant in the two in vitro conditions, suggesting that RPMI+10%BCS is more closely related to in vivo conditions than Spider medium. These data strongly support the notion that the transcriptional programs for filamentation are highly context dependent and that it is likely to be important to study regulators under multiple conditions.

In contrast to the highly context-dependent role of Rob1 during in vitro filamentation, the function of Rob1 during in vivo filamentation is much more consistent. Witchley et al. found that *rob1*ΔΔ mutants colonized the mouse GI tract predominantly as yeast (29). Consistent with these results, our group reported that Rob1 is one of the core regulators of filamentation during infection of subepithelial/mucosal tissue (12). In addition, *rob1*ΔΔ loss of function or deletion mutants form aberrant biofilms lacking hyphae in vivo (9). In this and previous work, we have found that *rob1*ΔΔ mutants can form filaments during OPC but they are less abundant and are not able to invade into submucosal tissue (11). Compared to the condition dependent roles of Rob1 in vitro, the relatively consistent phenotypes observed for *rob1*ΔΔ mutants in these distinct in vivo sites suggest that these niches share environment features that require Rob1 for filamentation.

The strain dependency of in vitro virulence phenotypes such as biofilm formation and filamentation has been well-established and is a continuing source of new insights into the pathobiology and diversity of *C. albicans* (10, 16, 21, 22, 26). A consistent outlier in surveys of the filamentation and biofilm phenotypes of *C. albicans* clinical isolates is the reference strain SC5314; it forms a high percentage of filaments in vitro and robust biofilms (22, 26). The genetic basis for the distinctions of SC5314 from other strains has been of significant interest. Clinical strains with reduced filamentation relative to SC5314 have been found to harbor loss of function mutations in *EFG1* (23) or increased activity of the filamentation repressor Nrg1 (5). As described above, SC5314 derived strains are heterozygous at the *ROB1* locus with a single SNP leading to to alleles with distinct functions. Surprisingly, all other *C. albicans* strains that we have examined are homozygous for the allele (*ROB1^946P^*) that has reduced filamentation and biofilm formation in vitro. These data suggest that the *ROB1^946S^* allele is a gain-of-function allele relative to the predominant *ROB1^946S^*allele. Further supporting that conclusion, introduction of *ROB1^946S^*into poorly filamenting strains homozygous for the *ROB1^946S^* allele increase their filamentation and converting an SC5314 strain to a homozygote of *ROB1^946S^* also further increases its ability to filament in relatively weak inducing conditions.

The presence of this *ROB1* gain of function allele is likely to contribute to the robust filamentation and biofilm phenotypes observed for SC5314, particularly in vitro. However, it is important not to overestimate the generality of these effects. First, there are other clinical isolates of *C. albicans* that have filamentation and biofilm phenotypes that are similar to SC5314 in vitro but are homozygous for the apparently less active *ROB1^946P^* allele (22). Second, the conversion of a poor filamenting clinical strain to the SC5314 heterozygous genotype at the *ROB1* locus improved its filamentation but it remained far less robust than SC5314. This observation is consistent with a similar experiment reported by Hirakawa et al (22) in which an *EFG1* loss of function allele was replaced by a functional allele in a poorly filamenting clinical isolate. In that case, filamentation improved but the levels remained well below that of SC5314. Consequently, it is likely that the phenotypic heterogeneity of these clinical isolates of *C. albicans* is due to genetic/genomic heterogeneity at multiple loci. Third, both alleles of *ROB1* support wild type levels of filamentation in mouse tissue and are indistinguishable in terms of virulence in this model. This indicates that the advantages or distinctions of the *ROB1^946S^* are dependent on environmental niche.

The distinctions between the functions of the two alleles in vivo are most apparent in settings that have features of the biofilm state. A consideration of these distinctions provides a possible model for the apparent low prevalence of the gain of function *ROB1^946S^* allele. The increased filamentation potential of the *ROB1^946S^* and the corresponding increased expression of inanimate surface biofilm promoting genes such as *HWP1* and *ALS3* are required for SC5314 to form a robust biofilm in a vascular catheter. However, the advantages of this allele disappear once *C. albicans* enters the blood stream to disseminate and infect other tissues. Similarly, both alleles support filamentation in oral tissue. The consequences of this filamentation, however, are starkly distinct. The *ROB1^946S^* allele promotes invasion into the submucosal compartment and triggers recruitment of inflammatory cells leading to reduced fungal burden. The strain bearing the *ROB1^946P^* allele, on the other hand, infects the oral tissue but is unable to penetrate or invade into the submucosal tissue. Accordingly, no immune cells are recruited to the site of infection. The OPC phenotype of the *ROB1^946P^* allele is very similar to the commensal phenotype described by Lemberg et al. for the commensal oral isolate 102 (5).

*C. albicans* is a commensal of the oral cavity and GI tract (1, 2, 3). Accordingly, the predominance of a commensal state-promoting *ROB1* allele in the population of sequenced clinical isolates is consistent with its niche as a commensal of human mucosae. The strain SC5314 is described in the literature as having been isolated from a patient with disseminated candidiasis (30). SC5314 is, therefore, not derived from a commensal niche but rather from a patient with disease. The presence of the *ROB1^946S^* that promotes invasion into the submucosal compartment raises the possibility that this allele may provide a competitive advantage over more commensal-oriented strains in the development of invasive disease. The rarity of the *ROB1^946S^*allele suggests that the ancestral allele is *ROB1^946P^* and that the SC5314 strain is a relatively new SNP. *C. tropicalis* also has a zinc cluster transcription factor that is homologous to Rob1. The *C. tropicalis* Rob1 homolog has a proline at the position corresponding to *C. albicans* Rob1946, further supporting the notion that this is the ancestral allele.

To date, we have been unable to gain detailed insights into the mechanism underlying the distinct phenotypes of the *ROB1^946S^* and *ROB1^946P^* alleles. As discussed above, the C-terminal region is a region where gain-of-function mutations frequently are found in zinc cluster TFs that affect fluconazole resistance. The C-terminal regions of some zinc finger TFs have been experimentally confirmed to be the activation domains of the factors and, thus, it is not surprising that a mutation such as those found in *ROB1* would lead to gain-of-function phenotypes. We have attempted to assess the relative activity of the two alleles in one-hybrid assays but have not been able to generate functional fusion proteins for this assay. Proline residues are well known to disrupt local and/or global domains in proteins and it is tempting to speculate that the decreased function of the *ROB1^946P^*may be related to that phenomenon. Future work will be required before we understand the molecular basis of the functional distinctions between these two *ROB1* alleles.

Finally, our data highlight the consequences of the heterozygous nature of the *C. albicans* genome on phenotypic diversity. Aneuploidy and loss-of-heterozygosity have well-established roles in driving the extensive phenotypic diversity of different *C. albicans* isolates (31). We show that a single amino acid change in Rob1 leads to significant in vivo and in vitro phenotypes. It is, therefore, likely that similar mutations alter function and that different combinations of heterozygous alleles within genes functioning in the same process or pathway also could lead to significant phenotypic diversity with respect to virulence or antifungal drug susceptibility (32).

## Materials and methods

### Strains, cultivation conditions and media

Genotypes and sources of *C. albicans* strains are summarized in Table S3. All *C. albicans* strains were precultured overnight in yeast peptone dextrose (YPD) medium at 30^0^C. Standard recipes were used to prepare synthetic drop-out media and YPD (14). RPMI medium was purchased and supplemented with bovine calf serum (10% or 1%, vol/vol). For filamentation assays, *C. albicans* strains were incubated overnight at 30^0^C in YPD media, harvested, and spotted on the plates at a concentration of 1 OD_600_. The plates were incubated at 37°C for three to five days prior to imaging. For liquid induction, the overnight cultures were diluted into RPMI + 10% BCS at a 1:50 ratio and incubated at 37^0^C for 4 h. Induced cells were fixed with 1% (v/v) formaldehyde. Fixed cells were then imaged using the Echo Rebel upright microscope with a 60x objective. The assays were conducted in triplicates on different days to confirm reproducibility.

### Strain construction

*Rob1 deletion mutants*. Both alleles of *ROB1* were deleted in clinical isolates using the transient CRISPR-Cas9 system (1). Briefly, cells were transformed with Cas9 and single *ROB1* guide RNA cassettes and the *rob1Δ::NAT1* repair template. The NAT1 repair template was amplified from the pCJN542 plasmid (generous gift from Dr. Aaron Mitchell, (2)) using primers with *ROB1* gene derived flanks (OL279, OL280, Table S4). The *ROB1* guide RNA was amplified by a split-join PCR using the OL1, OL2, 18.005, 18.008, 18.009 and 18.010 oligos. Transformants were selected on YPD media supplemented with 200 mg/mL nourseothricin and correct *rob1Δ::NAT1* repair template integration and absence of the *ROB1* ORF was checked by PCR using oligos derived from the *NAT1* gene, *ROB1* ORF and *ROB1 5’* and 3’ UTR (18.026, OL422 – OL425, Table S4).

ROB1 *allele swap*. *NAT1 knock-in* cassette carrying the respective SNP was made by three round PCR approach with first round primers (OL759, OL760, Supplementary methods) derived from 3’ end of the *ROB1* gene proximal to the 2902C/T SNP (corresponding to the 946S/P change in Rob1p) and *ROB1* 5’ UTR region. The first round product was then stitched to a *NAT1* cassette flanked with overlapping sequence in a second round PCR. The *NAT1* cassette with *ROB1* flanking sequence was amplified from pCJN542 using the OL757 and OL758 (Table S4). The stitched product was then amplified in a third round PCR using the flanking primers and used for transformation. Correct *knock-ins* were confirmed by PCR using *NAT1* and *ROB1* derived primers.

### In vitro biofilm growth and imaging

Biofilm growth was assayed as described in Glazier et al. (17). Briefly, strains were grown overnight at 30°C in liquid YPD with shaking, washed twice in phosphate-buffered saline (PBS), diluted to an OD_600_ of 0.5 in desired media. 200 μL of the suspension was dispensed into 6 wells of a 96-well plate (Corning Incorporated 96-well plate, catalog no. 3596). The cells were then incubated in a 37°C incubator for 90 minutes to allow adherence. Next, the media was removed, and the wells were washed once with PBS to remove non-adhered cells. The adherence density was measured by reading optical density using a SpectraMax plate reader. The biofilm inducing media was replaced and cells were further incubated at 37°C, without shaking. At 24 hours and 48 hours the media was aspirated; the cells were washed with PBS as above and the biofilm density was measured. Three biological replicates with 6 technical replicates per strain analyzed for each strain and the experiment was repeated independently 2-3 times. Differences between strains were analyzed by ANOVA and multiple comparison correction with statistical significance set at an adjusted P value <0.05. In vitro biofilm imaging and image processing was done as in (33) with minor modifications. Specifically, the strains were inoculated to an optical density at 600 nm (OD_600_) of 0.05 into prewarmed Spider media.

### In vivo filamentation assay

The inoculation and imaging of mice ear (female DBA/2 mice; 6-12 weeks) infections were carried out as described previously (12, 20). Acquired multiple Z stacks (minimum 15) were used to score the yeast vs filamentous ratio. A minimum of 100 cells from multiple fields were scored. Paired student’s *t* test with Welch’s correction (*P*>0.05) was used to define the statistical significance which was carried out using GraphPad prism software. Filament length of the *in vivo* samples were measured as described previously (12, 20). At least 50 cells per each strain from multiple fields were measured. A statistical significance was determined by Mann-Whitney U test (*P*>0.05).

### Gene expression analysis using quantitative-RT PCR

Strains were grown shaking overnight in liquid YPD at 30°C. Cells were back-diluted into fresh YPD, grown to mid-log phase for 4 hours, and then harvested. Total RNA was isolated with the MasterPure yeast RNA purification kit. The RNA was reverse transcribed using the iScript cDNA synthesis kit (170-8891; Bio-Rad), and quantitative reverse transcription-PCR (qPCR) was performed using IQ SyberGreen supermix (170-8882; Bio-Rad). *ACT1* expression was used as the normalization standard, and relative expression between strains and conditions were determined using the cycle threshold (ΔΔ*C_T_*) method. Experiments were performed in biological triplicate with technical triplicates, and the statistical significance was determined using Student’s *t* test or ANOVA followed by corrections for multiple comparisons with a significance limit of *P* < 0.05.

### RNA sequencing analysis of gene expression

The reference strain SN250 and the corresponding *rob1*ΔΔ mutant were subjected to hyphal induction conditions using either Spider medium or RPMI at 37°C for 4 hr. RNA was isolated as described above. Sequence reads were cleaned and adapter trimmed using trimmomatic-0.36 before mapping each sample individually to the *C. albicans* reference genome (SC5314 reference genome version A22 from the *Candida* Genome Database) with STAR2.5.2b. Raw read counts were obtained using featurecounts from the subread1.5.0p3 package and SC5314 gene annotations using only uniquely aligned reads (default) and including multi-mapping reads (-M). DESeq2-1.14.1 within R-3.3.2 (14-16) was used to perform data normalization and differential expression analysis with an adjusted p-value threshold of 0.05 on each set of raw expression measures.

### In silico analysis of *ROB1* alleles in sequenced *C. albicans* genomes

The analysis of *C. albicans* genome was performed as previous described (23). Whole genome sequences of 21 clinical isolates (NCBI BioProject ID: PRJNA193498), 43 clinical isolates from the OPC patients (NCBI BioProject ID: PRJNA257929), 8 clinical isolates from very low birth weight infant (VLBW) patient collection (25) and 182 isolates from various sources (NCBI BioProject ID: PRJNA432884) were obtained from previous studies (21-24) and aligned to the SC5314 reference genome version A22 from the *Candida* Genome Database (http://www.candidagenome.org) using the Burrows–Wheeler Alignment tool (version 0.7.17) with the BWA-MEM algorithm. SNP variants were called using the Genome Analysis Toolkit (GATK) (34) following GATK Best Practices. Polymorphisms were filtered using the GATK VariantFiltration tool by hard filters (QD<2.0, MQ<40.0, FS>60.0, MQRankSum<-12.5, ReadPosRankSum<-8.0). The variants of *ROB1* gene were manually checked in the Integrative Genomics Viewer (IGV).

### Rat model of *Candida albicans* vascular catheter infection

In vivo *C. albicans* biofilm formation was assessed using an external jugular-vein, rat-catheter infection model as previously described (35). Briefly, a 1x10^6^ cells/ml inoculum for each strain or strain combination was allowed to grow on an internal jugular catheter placed in a pathogen-free female rat (16-week old, 400 g) for 24 h. After this period, the catheter volumes were removed and the catheters were flushed with 0.9% NaCl. The biofilms were dislodged by sonication and vortexing. Viable cell counts were determined by dilution plating. Three replicates were performed for each strain. Differences in fungal burden were analyzed by ANOVA and correction for multiple comparisons with a significant difference defined as a P <0.05. Scanning electron microscopy of catheter biofilms. After a 24-hour biofilm formation phase, the devices were removed, sectioned to expose the intraluminal surface, and processed for SEM imaging. Briefly, one milliliter fixative (4% formaldehyde and 1% glutaraldehyde in PBS) was added to each catheter tube and tubes were fixed at 4 °C overnight. Catheters were then washed with PBS prior to incubation in 1% OsO4 for 30 min. Samples were then serially dehydrated in ethanol (30–100%). Critical point drying was used to completely dehydrate the samples prior to palladium–gold coating. Samples were imaged on a SEM LEO 1530, with Adobe Photoshop 2022 (v. 23.2.2) used for image compilation.

### Disseminated candidiasis model

Assessment of the fungal disease progression in a systemic candidiasis model was performed as described previously (11). Female CD-1 outbred mice (10 per group, 6 – 8 weeks old; Envigo) were inoculated by lateral tail vein injection with 1 x 10^6^ CFU of the indicated *C. albicans* strains and mice are monitored daily for clinical changes. Mice that demonstrated symptoms of severe diseases such as fur ruffling, difficulty with ambulation, abnormal posture, and/or failure to responds to surroundings was euthanized immediately. Disease progression was analyzed by Kaplan-Meier analysis and log rank (Mantel-Cox test, *P* < 0.05).

### Oropharyngeal candidiasis model

The immunosuppressed mouse model of OPC was employed, as previously described with some modifications (11). Male Balb/c mice were injected subcutaneously with cortisone acetate (300 mg/kg of body weight) on infection days: −1, 1, and 3. On the day of infection, the animals were sedated with ketamine and xylazine and a swab saturated with *C. albicans* strain SN250, *ROB1^946S^/rob1Δ* or *ROB1^946P^/rob1Δ* mutant (10^6^ cells per mL) was placed sublingually for 75 min. On post-infection day 5, the mice were sacrificed, and the tongues were harvested. For fungal burden studies, the harvested tongues were homogenized and plated for quantitative fungal burden (*n* = 5 per strain) or processed for histology. The log_10_-transformed fungal burden data for each experiment was analyzed by ANOVA with corrections for multiple comparisons to identify statistically significant differences between individual strains (adjusted *P <* 0.05).

## Acknowledgements

The authors thank Aaron Mitchell (Georgia) for assistance with biofilm imaging and review of drafts of the manuscript. This work was supported by the following grants from the NIH: F32AI26634 (VEG); R01AI073289 (DRA); R01AI141893 and R01AI081704 (RJB); R01DE026600 (SGF); R01AI133409 (DJK).

## Supplementary Material

**Figure S1. A.** SN250, *rob1-1*, *rob1-5*, and *rob1*ΔΔ strains were spotted on RPMI and RPMI+10% BCS plates and incubated for 3 days at either 30°C or 37°C before being photographed. The images are representative of two independent experiments. **B.** The filament lengths of the *rob1-1* and *rob1-5* strains were compared to SN250 24 hr post infection as described in materials and methods. There was no difference in length as determined by Mann-Whitney U test (P<0.05).

**Figure S2 A.** Schematic of *ROB1* allele swap construct. **B**.

**Table S1. RNA sequencing data comparing the expression profile of the *rob1*ΔΔ mutant to WT in Spider medium at 37°C after 4 hr of induction.**

**Table S2. RNA sequencing data comparing the expression profile of the *rob1*ΔΔ mutant to WT in RPMI medium at 37oC after 4 hr of induction.**

**Table S3. Table of strains**

**Table S4. Table of oligonucleotides.**

## References

1. Lopes JP, Lionakis MS. 2022. Pathogenesis and virulence of *Candida albicans*. Virulence 13: 89–121.

2. Vila T, Sultan AS, Montelongo-Jauregui D, Jabra-Rizk MA. 2020. Oral candidiasis: a disease of opportunity. J Fungi 16:15.

3. Pappas PG, Lionakis MS, Arendrup MC, Ostrosky-Zeichner L, Kullberg BJ. 2018. Invasive candidiasis. Nat Rev Dis Primers. 4:18026.

4. Sudbery PE. 2011. Growth of *Candida albicans* hyphae. Nat Rev Microbiol. 9:737–48.

5. Lemberg C, Martinez de San Vicente K, Fróis-Martins R, Altmeier S, Tran VDT, Mertens S, Amorim-Vaz S, Rai LS, d’Enfert C, Pagni M, Sanglard D, LeibundGut-Landmann S. 2022. Candida albicans commensalism in the oral mucosa is favoured by limited virulence and metabolic adaptation. PLoS Pathog. 18: e1010012.

6. Saville SP, Lazzell AL, Monteagudo C, Lopez-Ribot JL. 2003. Engineered control of cell morphology in vivo reveals distinct roles for yeast and filamentous forms of *Candida albicans* during infection. Eukaryot Cell. 2:1053–60.

7. Blankenship JR, Mitchell AP. 2006. How to build a biofilm. Curr Opin Microbiol. 9:588–94.

8. Ramage G, Borghi E, Rodrigues CF, Kean R, Williams C, Lopez-Ribot J. 2023. Our current clinical understanding of *Candida* biofilms: where are we two decades on? APMIS. doi: 10.1111/apm.13310.

9. Nobile CJ, Fox EP, Nett JE, Sorrells TR, Mitrovich QM, Hernday AD, Tuch BB, Andes DR, Johnson AD. 2012. A recently evolved transcriptional network controls biofilm development in *Candida albicans*. Cell. 148:126–138.

10. Do E, Cravener MV, Huang MY, May G, McManus CJ, Mitchell AP. 2022. Collaboration between antagonistic cell type regulators governs variation in the *Candida albicans* biofilm and hyphal gene expression network. mBio. 22:e0193722.

11. Solis NV, Wakade RS, Glazier VE, Ollinger TL, Wellington M, Mitchell AP, Filler SG, Krysan DJ. 2022. Systematic genetic interaction analysis identifies a transcription factor circuit required for oropharyngeal candidiasis. mBio. 13:e0344721.

12. Wakade RS, Ristow LC, Wellington M, Krysan DJ. 2023. Intravital imaging-based genetic screen reveals the transcriptional network governing *Candida albicans* filamentation during mammalian infection. Elife. 12:e85114.

13. Glazier VE, Murante T, Koselny K, Murante D, Esqueda M, Wall GA, Wellington M, Hung CY, Kumar A, Krysan DJ. 2018. Systematic complex haploinsufficiency-based genetic analysis of *Candida albicans* transcription factors: tools and applications to virulence-related phenotypes. G3 (Bethesda). 8:1299-1314.

14. Homann OR, Dea J, Noble SM, Johnson AD. 2009. A phenotypic profile of the *Candida albicans* regulatory network. PLoS Genet. 5:e1000783.

15. Noble SM, French S, Kohn LA, Chen V, Johnson AD. 2010. Systematic screens of a *Candida albicans* homozygous deletion library decouple morphogenetic switching and pathogenicity. Nat Genet. 42:590–598.

16. Huang MY, Woolford CA, May G, McManus CJ, Mitchell AP. 2019. Circuit diversification in a biofilm regulatory network. PLoS Pathog. 15:e1007787.

17. Glazier VE, Murante T, Murante D, Koselny K, Liu Y, Kim D, Koo H, Krysan DJ. 2017. Genetic analysis of the *Candida albicans* biofilm transcription factor network using simple and complex haploinsufficiency. PLoS Genet. 13:e1006948.

18. Xu W, Solis NV, Ehrlich RL, Woolford CA, Filler SG, Mitchell AP. 2015. Activation and alliance of regulatory pathways in *C. albicans* during mammalian infection. PLoS Biol. 13:e1002076.

19. Cravener MV, Do E, May G, Zarnowski R, Andes DR, McManus CJ, Mitchell AP. 2023. Reinforcement amid genetic diversity in the *Candida albicans* biofilm regulatory network. PLoS Pathog. 19:e1011109.

20. Wakade RS, Krysan DJ, Wellington M. 2022. Use of in vivo imaging to screen for morphogenesis phenotypes in *Candida albicans* mutant strains during active infection in a mammalian host. J Vis Exp. 212;188. doi: 10.3791/64258.

21. Ford CB, Funt JM, Abbey D, Issi L, Guiducci C, Martinez DA, Delorey T, Li BY, White TC, Cuomo C, Rao RP, Berman J, Thompson DA, Regev A. 2015. The evolution of drug resistance in clinical isolates of *Candida albicans*. Elife. 4:e00662.

22. Hirakawa MP, Martinez DA, Sakthikumar S, Anderson MZ, Berlin A, Gujja S, Zeng Q, Zisson E, Wang JM, Greenberg JM, Berman J, Bennett RJ, Cuomo CA. 2015. Genetic and phenotypic intra-species variation in *Candida albicans*. Genome Res. 25:413–425.

23. Liang SH, Anderson MZ, Hirakawa MP, Wang JM, Frazer C, Alaalm LM, Thomson GJ, Ene IV, Bennett RJ. 2019. Hemizygosity enables a mutational transition governing fungal virulence and commensalism. Cell Host Microbe. 25:418–431.

24. Ropars J, Maufrais C, Diogo D, Marcet-Houben M, Perin A, Sertour N, Mosca K, Permal E, Laval G, Bouchier C, Ma L, Schwartz K, Voelz K, May RC, Poulain J, Battail C, Wincker P, Borman AM, Chowdhary A, Fan S, Kim SH, Le Pape P, Romeo O, Shin JH, Gabaldon T, Sherlock G, Bougnoux ME, d’Enfert C. 2018. Gene flow contributes to diversification of the major fungal pathogen Candida albicans. Nat Commun. 9:2253.

25. Bliss JM, Basavegowda KP, Watson WJ, Sheikh AU, Ryan RM. 2008. Vertical and horizontal transmission of *Candida albicans* in very low birth weight infants using DNA fingerprinting techniques. Pediatr Infect Dis J. 27:231–5.

26. Anderson FM, Visser ND, Amses KR, Hodgins-Davis A, Weber AM, Metzner KM, McFadden MJ, Mills RE, O’Meara MJ, James TY, O’Meara TR. 2023. Candida albicans selection for human commensalism results in a substantial within-host diversity without decreasing fitness for invasive disease. PLoS Biol. 21:e3001822.

27. MacPherson S, Larochelle M, Turcotte B. 2006. A fungal family of transcriptional regulators: the zinc cluster proteins. Microbiol Mol Biol Rev. 70:583–604.

28. Sasse C, Dunkel N, Schäfer T, Schneider S, Dierolf F, Ohlsen K, Morschhäuser J. 2011. The stepwise acquisition of fluconazole resistance mutations causes a gradual loss of fitness in *Candida albicans*. Mol Microbiol. 86:539–56.

29. Witchley JN, Penumetcha P, Abon NV, Woolford CA, Mitchell AP, Noble SM. 2019. *Candida albicans* morphogenesis programs control the balance between gut commensalism and invasive infection. Cell Host Microbe. 25:432–443.

30. Gillum AM, Tsay EY, Kirsch DR. 1984. Isolation of the *Candida albicans* gene for orotidine-5’-phosphate decarboxylase by complementation of *S. cerevisiae ura3* and *E. coli* pyrF mutations. Mol Gen Genet 198:179–182.

31. Ene IV, Bennett RJ, Anderson MZ. 2019. Mechanisms of genome evolution in *Candida albicans*. Curr Opin Microbiol. 52:47–54.

32. Liang SH, Bennett RJ. 2019. The impact of gene dosage and heterozygosity on the diploid pathobiont *Candida albicans*. J Fungi (Basel). 6:10.

33. Sharma A, Solis NV, Huang MY, Lanni F, Filler SG, Mitchell AP. 2023. Hgc1 Independence of Biofilm Hyphae in Candida albicans. mBio. 14:e0349822.

34. McKenna A, Hanna M, Banks E, Sivachenko A, Cibulskis K, Kernytsky A, Garimella K, Altshuler D, Gabriel S, Daly M, DePristo MA. 2010. The Genome Analysis Toolkit: a MapReduce framework for analyzing next-generation DNA sequencing data. Genome Res. 20:1297–303.

35. Nett JE, Marchillo K, Andes DR. 2012. Modeling of fungal biofilms using a rat central vein catheter. Methods Mol Biol. 845:547–556.

